# A full-length single nuclei transcriptomic atlas of human skeletal muscle insulin resistance

**DOI:** 10.64898/2026.02.26.708250

**Authors:** Katie L Whytock, Adeline Divoux, James Vazquez, Meghan Hopf, Mark R Viggars, Miguel A Gutierrez-Monreal, Courtney H. Ruggiero, Felix R. Jimenez-Rondan, Francielly Morena, Polina Krassovskaia, Nicholas T. Broskey, Yifei Sun, Martin J Walsh, Robert J Cousins, Joseph A Houmard, Lauren M Sparks, Bret H Goodpaster

**Affiliations:** Translational Research Institute, AdventHealth, Orlando, FL, USA; Department of Physiology and Aging, College of Medicine, University of Florida, Gainesville, FL, USA; Center for Nutritional Sciences, Food Science and Human Nutrition Department, College of Agricultural and Life Sciences, University of Florida, Gainesville, FL, USA; Department of Cancer Biology, Wake Forest University, Winston Salem, North Carolina, USA; Department of Kinesiology, Human Performance Laboratory, Easy Carolina Diabetes and Obesity Institute, East Carolina University, Greenville, North Carolina, USA; Icahn School of Medicine at Mount Sinai, New York, USA

## Abstract

Skeletal muscle (SkM) insulin resistance is a central defect in T2D, yet cell specific molecular determinants remain incompletely understood. Here, we integrate full-length single-nucleus transcriptomics with gold-standard stable isotope-labeled hyperinsulinemic-euglycemic clamps to generate a nucleus-resolved transcriptomic atlas of SkM insulin resistance. We identify previously unrecognized myonuclear populations whose proportions associate with insulin sensitivity across independent cohorts, revealing MYH7B+ myonuclei are metabolically favorable over EGF+ myonuclei. Modeling transcriptional variation against tracer-derived glucose disposal uncovers highly nucleus-specific molecular programs that are obscured when using surrogate fasting indices. Mechanistically, we identify zinc transporter ZIP14 as a positive regulator of insulin-stimulated glucose uptake and implicate EGF signaling in impaired branched-chain amino acid catabolism and inflammatory cross-talk within the SkM niche. Together, these findings redefine SkM insulin resistance as a multicellular, nucleus-resolved process and highlight new cell type specific targets for metabolic intervention.

## Introduction

Type 2 Diabetes (T2D) is estimated to affect over 500 million people worldwide and vastly increases the risk of cardiovascular complications^1^. Skeletal muscle (SkM) is one of the primary sites of glucose uptake following insulin stimulation^2–4^. While the pathogenesis of T2D is heterogeneous^5^, defects in SkM insulin responsiveness are considered one of the primary aberrations in the development of T2D^6–8^. Among the tissues affected with T2D, SkM is recognized for its ability to adapt or mal-adapt to different environmental stimuli such as exercise, nutrition and pharmaceuticals^9,10^ making it a significant therapeutic target.

With the advent of transcriptomics technologies, research has shown dysregulated transcriptional profiles in SkM insulin resistance related to glucose metabolism, insulin signaling, inflammation and amino acid metabolism^11,12^. With these rich datasets, researchers are equipped to identify novel strategies to improve insulin sensitivity including drug repurposing efforts^13^. A significant limitation of these prior studies, however, is the restriction to whole SkM homogenates to perform bulk transcriptomics.

SkM is comprised of; myofibers, endothelial cells (EC), smooth muscle cells (SMC), fibro-adipogenic progenitors (FAP), muscle satellite cells (MuSC) and immune cells that all play a role in SkM metabolic health^14^. Effective insulin-stimulated SkM glucose uptake is governed by; 1) insulin-stimulated activation of terminal arteriole ECs that activate vasodilation of SMC to perfuse microvascular units, 2) transport of substrates across capillary EC and 3) stimulation of myofiber insulin signaling resulting in glucose uptake^15–20^. Each of these steps has the potential to be dysregulated by pro-inflammatory cytokines released from localized immune cells in the SKM niche^14^. Further adding to this complexity are myofibers being uniquely characterized by their long length >2-3cm and multi-nucleated syncytia. Myonuclei are dispersed along the myofiber and are responsible for the mRNA transcription of a finite volume of cytoplasm termed the myonuclear domain^21–23^. Recent snRNA-seq analyses of SkM shows remarkable transcriptional heterogeneity among myonuclei^24,25^ raising the possibility that distinct myonuclear populations contribute differentially to SkM insulin resistance.

Advancements in single cell (sc) and single nuclei (sn) RNA-seq have permitted cell/nuclei-type specific transcriptional profiling related to aging and different metabolic disease states^26–30^. To date, all sc/sn RNA-seq research in SkM have been performed with technology that only permits 3’ or 5’ amplification of the gene body which results in lower gene coverage per cell. We leveraged our recent advancements in full-length sc/sn RNAseq profiling with the ICELL8 system^31,32^ which amplifies both 3’ and 5’ ends to perform, to our knowledge, full-length snRNA-seq profiling in human SkM for the first time. A hyperinsulinemic-euglycemic (HE) clamp with stable isotope glucose tracers is considered the gold standard method for quantifying SkM insulin sensitivity because it directly quantifies insulin-stimulated glucose disposal. Our aim therefore was to combine gold standard *in vivo* phenotyping of SkM insulin sensitivity with high dimensional transcriptional profiling to elucidate nuclei-specific transcriptional profiles of human SkM insulin resistance.

## Results

### Phenotypical characteristics

To explore the effects insulin resistance on SkM transcriptome at a single nuclei level, we performed full-length snRNA-Seq on the vastus lateralis of older (> 60 years old) individuals with obesity with T2D (T2D) (n = 9) and without T2D (CON) (n = 9) (**Figure 1A**) that had similar age, BMI and aerobic capacity (VO_2peak_) (Table 1). Both CON and T2D had comparable cholesterol profiles (**Table 1**). Overall fat mass and body fat % were also similar between groups; however, T2D had greater VAT mass and lean mass (*P* < 0.05). As expected, individuals with T2D had impaired glycemic control evident by elevated fasting glucose, fasting insulin, HOMA2-IR and HbA1C (%).

**Figure 1.**
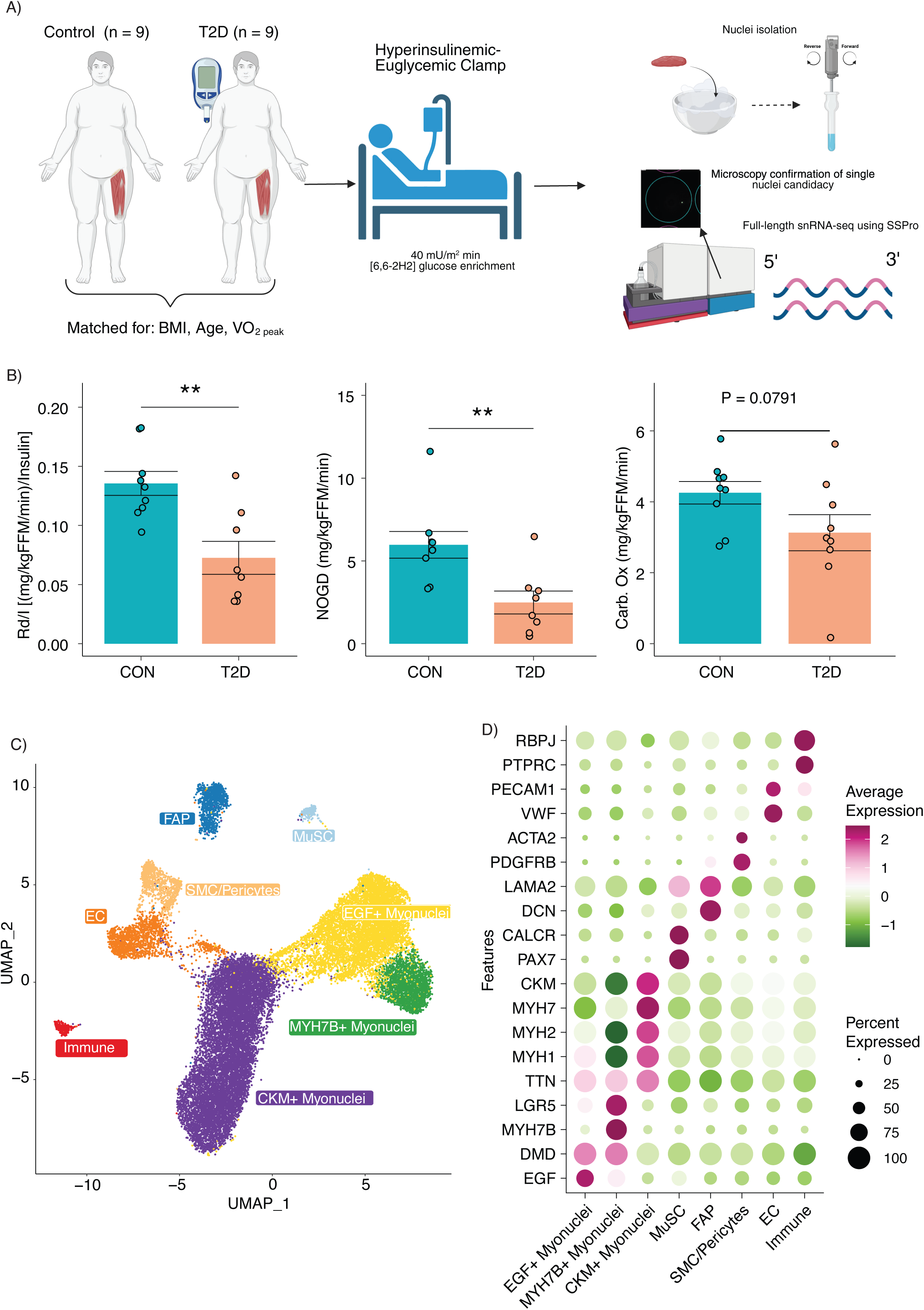
A full-length single nuclei transcriptomic atlas of human skeletal muscle insulin resistance. A schematic of participants characteristics, deep phenotyping and library preparation workflow (A). Differences between CON and T2D for skeletal muscle insulin sensitivity indices derived from a hyperinsulinemic-euglycemic clamp with stable glucose isotope tracers (B). A UMAP of 25,303 nuclei from 18 human skeletal muscle samples clustered into different nuclei populations (C). A dotplot of differentially expressed genes and cell marker genes for each nuclei population (D). *, *P* < 0.05.

**Table 1:** Participants Characteristics.

| Characteristic | CON (n = 9) | T2D (n = 9) | P Value |
| --- | --- | --- | --- |
| Sex (F/M) | (8/1) | (8/1) |  |
| Age (years) | 67 ± 4 | 66 ± 3 | 0.606 |
| BMI (kg/m<sup>2</sup>) | 34.47 ± 3.13 | 36.08 ± 3.51 | 0.321 |
| Weight (kg) | 88.30 ± 5.68 | 97.81 ± 11.48 | **0.0407** |
| WC (cm) | 109.47 ± 5.22 | 112.67 ± 9.55 | 0.392 |
| DBP | 73 ± 9.64 | 72.56 ± 11.90 | 0.932 |
| SBP | 129.00 ± 16.02 | 134.89 ± 9.39 | 0.355 |
| Cholesterol (mmol/L) | 201.89 ± 41.78 | 186.56 ± 45.70 | 0.48 |
| HDL (mmol/L) | 59.22 ± 20.03 | 53.00 ± 13.87 | 0.455 |
| LDL (mmol/L) | 113.89 ± 40.33 | 100.78 ± 38 | 0.488 |
| Triglycerides (mmo/L) | 143.22 ± 73.80 | 73.80 ± 66.27 | 0.48 |
| VLDL (mmol/L) | 28.78 ± 14.83 | 32.78 ± 13.15 | 0.4 |
| Abdomen AT (kg) - MRI | 21.81 ± 2.07 | 24.63 ± 5.31 | 0.209 |
| Abdomen SAT (kg) - MRI | 15.84 ± 1.88 | 15.66 ± 3.76 | 0.914 |
| Abdomen VAT (kg) - MRI | 5.98 ± 1.19 | 8.96 ± 1.98 | **0.0035** |
| FFM (kg) - DXA | 45.93 ± 4.95 | 53.19 ± 5.07 | **0.00731** |
| Fat Mass (kg) - DXA | 42.95 ± 4.79 | 44.72 ± 8.69 | 0.598 |
| Lean Mass (kg) - DXA | 43.60 ± 4.83 | 50.39 ± 4.73 | **0.00829** |
| Body Fat (%) – DXA | 48.33 ± 4.56 | 45.41 ± 4.28 | 0.18 |
| VO<sub>2</sub> peak (ml/kgFFM/min) | 29.28 ± 5.28 | 28.80 ± 4.54 | 0.859 |
| Fasting Glucose (mg/dL) | 97.22 ± 4.06 | 111.78 ± 17.29 | **0.0257** |
| Fasting insulin (mIU/mL) | 8.77 ± 4.59 | 19.96 ± 9.56 | **0.0146** |
| HOMA2-IR | 1.15 ± 0.59 | 2.65 ± 1.2 | **0.0114** |
| HbA1c (%) | 5.70 ± 0.34 | 6.72 ± 0.87 | **0.00477** |

HE clamp with stable glucose isotope tracers is the gold standard method for quantifying SkM insulin sensitivity and is quantified as rate of glucose disposal (mg/kgFFM/min) divided by plasma insulin (Rd/I) during steady state. Individuals with T2D had impaired Rd/I compared to CON (**Figure 1B**) and there was a large variability within the data allowing us to conduct downstream association analyses with these variables. When glucose enters SkM it is either stored as glycogen or oxidized. By incorporating indirect calorimetry into the HE clamps, we also calculated insulin-stimulated glycogen storage, referred to as non-oxidative glucose disposal (NOGD), and carbohydrate oxidation. In line with previous findings the main aberration with SkM insulin resistance and T2D was a reduced NOGD^33,34^ (**Figure 1B**) with only a trend (*P* = 0.0791) for a reduction in carbohydrate oxidation (**Figure 1B**).

### Single nuclei transcriptomics atlas of SkM insulin resistance

Our single nuclei transcriptomic atlas of SkM insulin resistance was composed of 25,303 nuclei that detected on average 4962 genes per nuclei. Initial clustering resolved 8 distinct clusters that were annotated with differentially expressed genes (**Supplementary Table 1**) and known marker genes (**Figure 1C-D**). Among nuclei types were mononuclear cells; muscle satellite cells (MuSC) (*PAX7*), fibro-adipogenic progenitors (FAP) (*DCN*), Immune (*PTPRC*), EC (*VWF*) and SMC/Pericytes (*PDGFRB*). Due to the myofibers being syncytia, the majority (84%) of nuclei profiled were myonuclei (*TTN*), in line with previous data^29^. Each participant had a representation of every nuclei-type profiled (**Supplementary Figure 1A**). Quality control metrics were comparable between nuclei types and samples (**Supplementary Figure 1B-C**).

#### Novel Myonuclei populations

The myonuclei populations did not originally cluster based on traditional myosin heavy chain expression (MYH7/Type I; MYH2/Type IIa; MYH1/Type IIx) and instead were marked by high expression of either *EGF*, *MYH7B* or *CKM*. CKM+ myonuclei exhibited high expression of *MYH1/2/7* and when this cluster was stratified based on traditional myosin heavy chain expression alone, the nuclei could be classified into Type I, IIa, IIx and hybrid myonuclei (**Supplementary Figure 2A**). The proportions of Type I, IIa, IIx and hybrid myonuclei showed a similar distribution pattern to myofibers quantified with immunohistochemistry (**Supplementary Figure 2B**), except snRNA-seq showed lower Type I proportion irrespective of T2D diagnosis which may be explained by differences in measuring RNA vs protein or the concept that multiple nuclei exist within a fiber.

Myosin heavy chains are just singular markers and when the whole transcriptome was taken into consideration, myonuclei clustered based on different genes and pathways. EGF+ myonuclei were characterized by genes related to action potential, intracellular signaling and contraction while MYH7B+ myonuclei were characterized by genes related to cation transport (**Figure 2A**). In contrast, CKM+ myonuclei had an upregulation of genes related to oxidative phosphorylation (**Figure 2A**).

**Figure 2.**
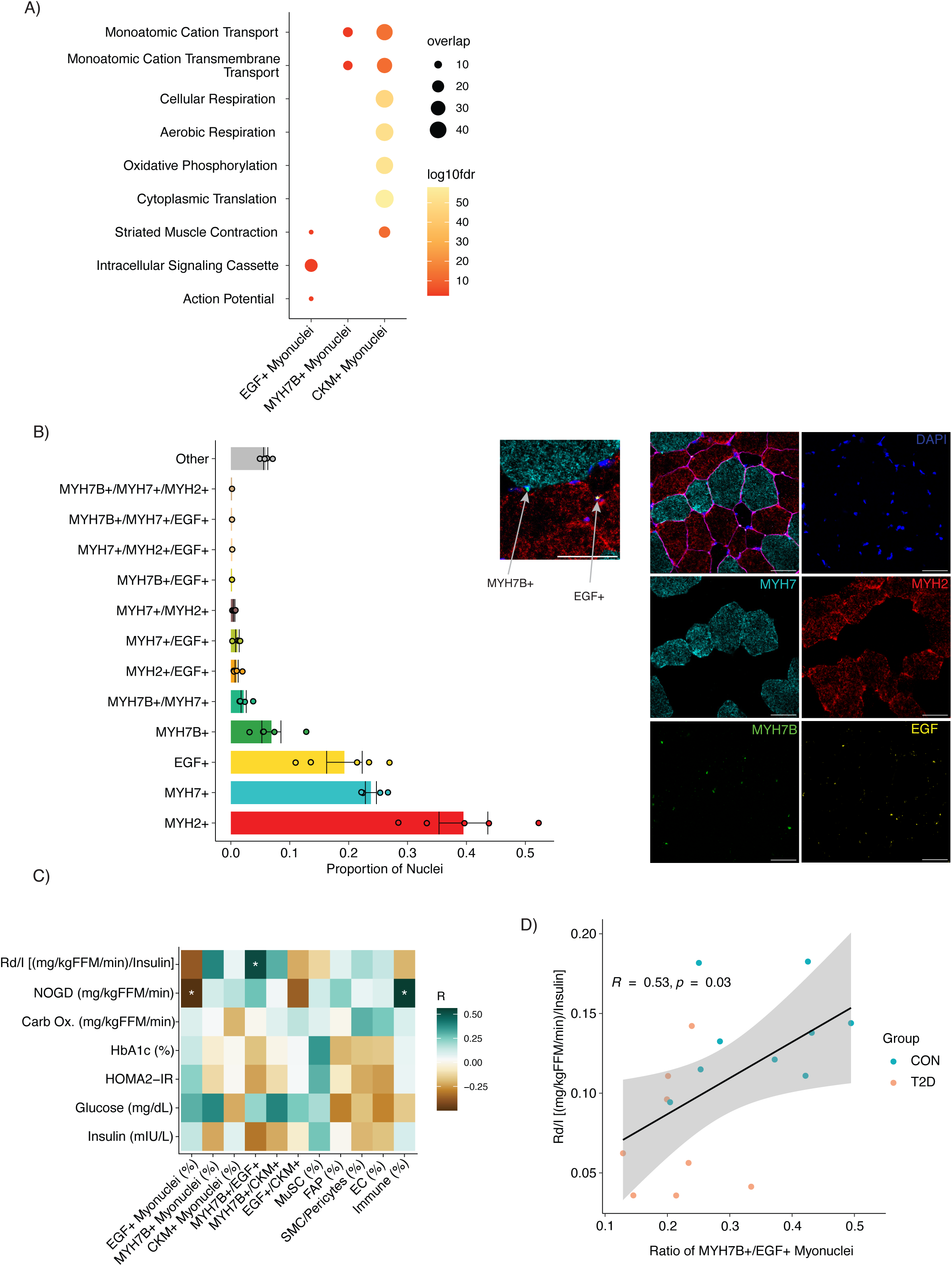
Novel myonuclei populations are associated with skeletal muscle insulin sensitivity. Pathway analyses of upregulated genes of different myonuclei populations (A). Validation and quantification of novel myonuclei populations with RNAscope (B). Correlations matrix between proportion of nuclei populations and glycemic control indices including variables from hyperinsulinemic-euglycemic clamp (C). The ratio of MYH7B+/EGF+ myonuclei correlates positively with Rd/I (D). *, *P* < 0.05.

Alternative splicing events prevent MYH7B protein production in mammalian skeletal muscle^35^. This was supported in our data with *MYH7B* gene expression being driven by non-protein coding transcripts (MYH7B- 209 and MYH7B-207) (**Supplementary Figure 2C**). We therefore validated these new myonuclei populations (MYH7B+ and EGF+) with RNAscope, while simultaneously probing for conventional myosin heavy chains *MYH2* and *MYH7* that together represented the CKM+ myonuclei population. Both *MYH2* and *MYH7* staining was diffuse throughout the myofibers whereas *MYH7B* and *EGF* staining was more punctate and mainly localized to the myonuclei (**Figure 2B**). While most myonuclei were MYH2+ or MYH7+; a substantial fraction were EGF+ or MYH7B+ (**Figure 2B**). Co-expression of *MYH7B* occurred primarily with *MYH7* whereas EGF showed equal co-expression with *MYH2* and *MYH7* (**Figure 2B**). There was a limited proportion of myonuclei with co-expression of 3 target genes (**Figure 2B**). Nuclei classified as other represent non-myonuclei proportions present within the muscle tissue e.g. EC (**Figure 2B**). This data confirms myonuclei transcriptional heterogeneity exists beyond traditional myosin heavy chains, in line with previous work using isolated myofibers^36,37^

#### Nuclei Type Proportions

There were no significant differences in nuclei proportions between CON and T2D (**Supplementary Figure 2D**) except for a trend towards T2D having greater EGF+ myonuclei (*P* = 0.063). Nuclei proportions did, however, correlate with glycemic control indices (**Figure 2C**). The proportion of EGF+ myonuclei and the proportion of immune cells negatively and positively correlated with NOGD respectively (**Figure 2C**). The ratio of MYH7B+ to EGF+ myonuclei positively correlated with Rd/I (**Figure 2D**), suggesting MYH7B+ rather than EGF+ myonuclei exert a positive metabolic influence on SkM insulin sensitivity. We next deconvoluted published bulk RNA-seq datasets to identify whether EGF+ and MYH7B+ nuclei proportions also correlate with glycemic control indices in other datasets. In the CALERIE study, Matsuda insulin sensitivity index (ISI) was calculated from an oral glucose tolerance test^38,39^. At baseline the ratio of MYH7B+/EGF+ nuclei positively correlated with Matsuda ISI (*P* < 0.05) (**Supplementary Figure 2E**) and this was primarily driven by differences in MYH7B+ proportion. Following 24 months of intervention, there was a positive correlation between the % change in MYH7B+ proportion and the % change in Matsuda ISI (*P* < 0.05) (**Supplementary Figure 2E**).

As our cohort were all older individuals with obesity and mainly female, we next investigated whether MYH7B+/EGF+ myonuclei ratio were altered with age, BMI or sex in publicly available bulk RNA-seq data sets. MYH7B+/EGF+ myonuclei ratio was significantly greater in older (60-85 years) males compared to younger (20-35 years) males (*P* < 0.05) that had similar levels of leanness and activity levels in a study by Trim et al.,^40^ (GSE175495) (**Supplementary Figure 2F**). This finding was replicated in a study of older (60-75 years) and younger (18-35 years) individuals without obesity that underwent an insulin resistance inducing bed-rest intervention (5 days)^41^ (GSE113165). At baseline the older group had a greater ratio of MYH7B/EGF+ myonuclei compared to the younger group (*P* < 0.01); however, MYH7B/EGF+ myonuclei ratio decreased following bed-rest in the older group (*P* < 0.05) to similar levels as the younger group (**Supplementary Figure 2G**). Using the GTEx data which was obtained with from the *gastrocnemius* rather than the *vastus lateralis*, we observed greater MYH7B+/EGF+ myonuclei ratio in individuals aged between 50-59 years compared to individuals aged between 30-39 or 40-49 years (*P* < 0.05). This greater MYH7B+/EGF+ myonuclei ratio was retained in the 60-69 year age bracket before decreasing in individuals between 70-79 years (*P* < 0.05) (**Supplementary Figure 2H**). There were was also a main effect for sex (*P* < 0.05) with males having a higher ratio of MYH7B+/EGF+ myonuclei (**Supplementary Figure 2H**). Lastly, in a study comparing males of different BMI^42^, there was no difference in MYH7B+/EGF+ myonuclei between lean males and males with overweight/obesity (**Supplementary Figure 2I**).

Together, this data suggests that MYH7B+/EGF+ myonuclei ratio is higher in males and increases with healthy aging independently of BMI. This favorable MYH7B+/EGF+ myonuclei ratio is lowered in individuals with insulin resistance and decreases with interventions that induce insulin resistance.

### Nuclei-specific transcriptional differences with T2D and insulin resistance

#### Single Nuclei Transcriptional Profiles with T2D

Pseudobulk differential gene expression analyses between T2D and CON revealed the greatest transcriptional differences in non-myonuclei populations (**Supplemental Figure 3A**, **Supplementary Table 4**), however these transcriptional differences did not result in any significantly enriched pathways (FDR < 0.05). The low number of DEGs and pathways between CON and T2D SkM likely reflects the complex, multifactorial nature of T2D, wherein diverse and overlapping factors such as hyperglycemia, insulin resistance and medication use can exert heterogenous and uncoordinated effects on the transcriptome. Similar conclusions were drawn when investigating the SkM proteome of individuals with T2D compared to individuals with normal glucose tolerance, where there was a minimal number of differentially abundant proteins^43^. This supports the approach of modelling molecular markers to key continuous variables that are directly linked to the impacted tissue e.g. SkM insulin sensitivity rather than comparing discrete groups.

#### Single Nuclei Transcriptional Profiles with SkM insulin resistance

SkM insulin sensitivity is a primary aberration in the development of T2D^6^ and is quantified most accurately by a HE clamp with stable isotope tracers to derive Rd/I. Rd/I had a 44% CV among individuals showing high variability for association analyses and therefore was modelled against nuclei specific transcriptional signatures. Correlation analyses between Rd/I and gene expression revealed remarkable nuclei-specific signatures (**Figure 3A, Supplementary Table 3**). Out of the total 4867 total genes correlating with Rd/I across all nuclei sub-types, only 780 genes overlapped with more than one cell-type. This suggests that different cell types within SkM tissue have different molecular profiles related to insulin resistance.

**Figure 3.**
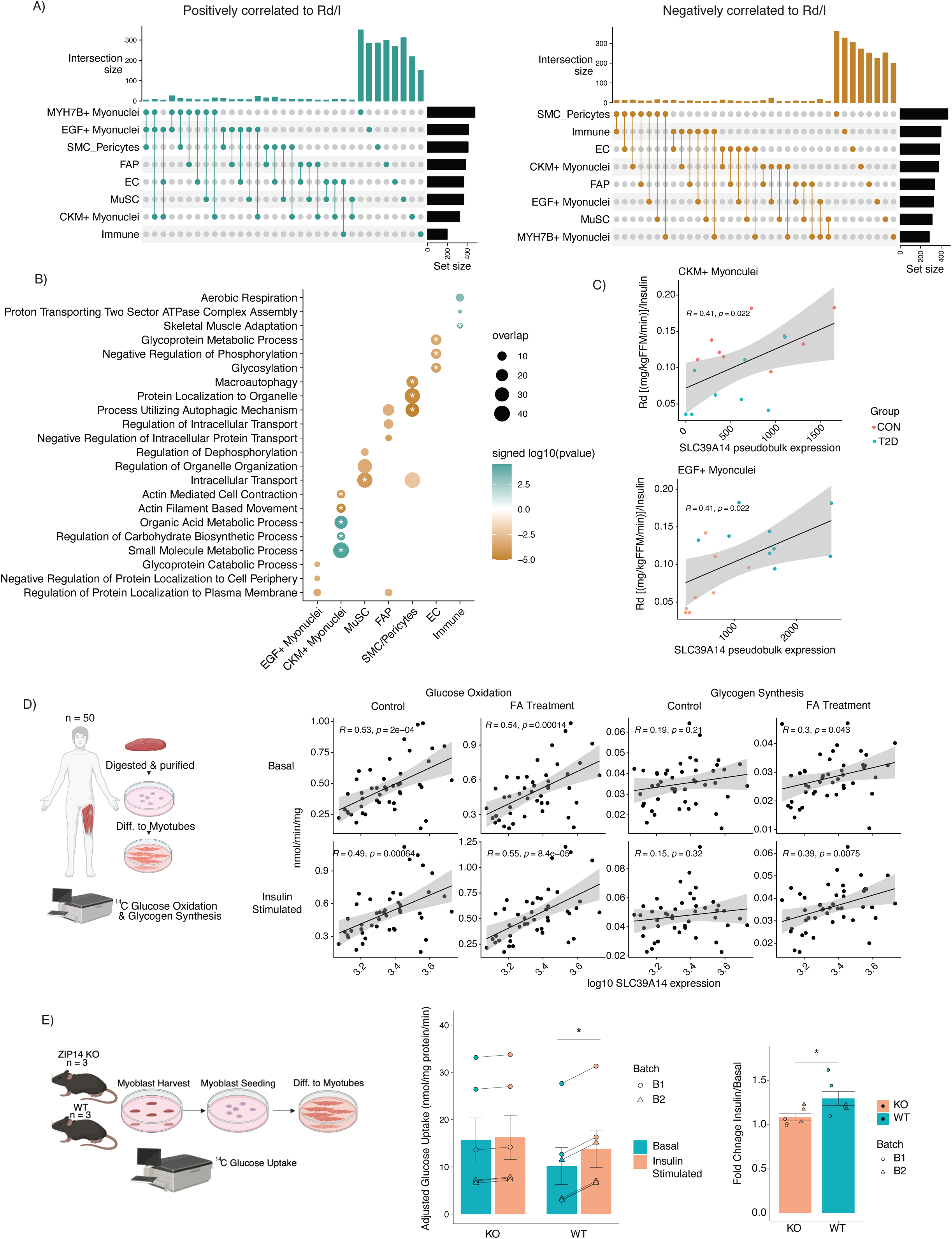
Nuclei specific transcriptional signatures associated with SkM insulin resistance. Upset plots highlighting the overlap of genes correlated to Rd/I across the different nuclei-types (A). Pathway analyses of genes positively and negatively associated with insulin resistance in each nuclei type (B) * in dotplot is FDR < 0.2. *SLC39A14* gene expression positively correlated with Rd/I in both CKM+ Myonuclei and EGF+ Myonuclei (C). Correlations between SLC39A14 gene expression and glucose uptake indices in primary human differentiated myotubes (D). Basal and insulin-stimulated glucose uptake in differentiated myotubes from ZIP14 KO and WT mice (E). *, *P* < 0.05.

Pathway analyses revealed that genes related to carbohydrate related processes were positively correlated to Rd/I in CKM+ myonuclei only, whereas genes related to protein localization to the plasma membrane were negatively correlated to Rd/I in the EGF+ myonuclei (**Figure 3B, Supplementary Table 4**). MYH7B+ myonuclei did not have any distinct pathways positively or negatively correlated to Rd/I indicating that unlike the proportion of MYH7B+ myonuclei, differences in the transcriptome of MYH7B+ myonuclei may not be directly linked to SkM insulin sensitivity. SMC/Pericytes had autophagy and protein localization related genes negatively correlated to Rd/I, whereas EC had genes related to glycosylation and growth negatively correlated to Rd/I (**Figure 3B, Supplementary Table 4**).

#### Single Nuclei Transcriptional Profiles with HOMA2-IR

Whilst Rd/I is the gold standard measure of SkM insulin sensitivity, it is not always an accessible measure in research populations. Therefore, previous work has modeled SkM gene expression to HOMA2-IR^12^ which is derived from only fasting glucose and insulin^44^. To compare, we also correlated nuclei-specific gene expression to HOMA2-IR (**Supplementary Table 5**). Pathway analyses revealed genes related to protein catabolism were upregulated with HOMA2-IR among several nuclei-types indicative of a lack of nuclei specificity when modelling to HOMA2-IR (**Supplementary Table 6**). Previous findings also highlighted BCAA catabolism genes related to HOMA2-IR when gene expression was measured in whole muscle homogenates^12^. We next compared the overlap of genes correlated to HOMA2-IR and genes correlated to Rd/I on a nuclei-specific basis. For each nuclei-type there was minimal overlap between genes that correlated with improved insulin sensitivity (positive correlation to Rd/I and negative correlation to HOMA2-IR) and with genes that correlated with reduced insulin sensitivity (negative correlation to Rd/I and positive correlation to HOMA2-IR) (**Supplementary Figure 3B**). This suggests SkM transcriptional signatures should be modelled to gold standard SkM insulin sensitivity measures to detect molecular features in SkM related to impaired insulin-stimulated glucose uptake. The intersect between genes that correlate with Rd/I and HOMA2-IR can be considered important genes for monitoring improvements in SkM when only HOMA2-IR indices are available.

##### The impact of ZIP14 on insulin-stimulated glucose uptake

*SLC39A14* is a part of the small molecule metabolic process pathway that positively correlated to Rd/I in both CKM+ and EGF+ myonuclei populations (**Figure 3C**). *SLC39A14* encodes zinc transporter ZIP14 which has previously been implicated in regulating hepatocyte glucose metabolism^45^ and skeletal muscle myogenesis through suppression of metabolic endotoxemia^46^. We sought to validate the role of ZIP14 in SkM insulin-stimulated glucose uptake using two independent datasets. We recently published the effects of habitual exercise training on basal, and insulin-stimulated glucose oxidation and glycogen synthesis rates with and without a insulin resistance inducing fatty-acid (FA) treatment on primary human skeletal muscle stem cells differentiated to myotubes *in vitro*^47^. Combining this functional data-set of 50 individual samples with concurrent RNA-seq, we probed how *SLC39A14* gene expression correlated to these glucose uptake metrics. *SLC39A14* correlated positively with all glucose oxidation metrics (*P <* 0.001) irrespective of FA treatment or insulin stimulation (**Figure 3D**). Conversely, *SLC39A14* correlated positively with glycogen synthesis rates only when treated with FA during both basal conditions and following insulin stimulation (*P <* 0.05) (**Figure 3D**), suggesting *SLC39A14* is more beneficial for glycogen synthesis in insulin-resistance inducing conditions. Lastly, skeletal muscle was harvested from WT mice and ZIP14 KO mice and myoblasts were isolated and differentiated into myotubes and tested for basal and insulin-stimulated glucose uptake *in vitro*. *SLC39A14/ZIP14 expression* was only present in the WT cells (**Supplementary Figure 3C**) and both cell lines differentiated into myotubes as shown by changes in expression of Myod, Myh2 and Myh3 (**Supplementary Figure 3D**). There was a significant increase in glucose uptake with insulin stimulation in WT myotubes (*P* < 0.05) and not in the KO myotubes (**Figure 3E**). This resulted in WT myotubes having a significantly greater fold change in glucose uptake from basal to insulin stimulation compared to myotubes from the ZIP14 KO mice (*P* < 0.05) (**Figure 3E**). Together this data suggests *SLC39A14*/ZIP14 aids in insulin-stimulated glucose uptake and may exert more influence in an insulin-resistance microenvironment.

### EGF cell communication with SkM insulin resistance

EGF+ myonuclei proportions showed unfavorable associations with metabolic health and had distinct transcriptional profiles related to SkM inulin sensitivity. As EGF is a ligand with an established receptor, EGFR, we next evaluated how EGF cellular cross-talk occurs within the SkM niche and how it is affected with SkM insulin resistance. This was achieved with two complementary bioinformatic approaches that A) predicted cell-to-cell communication based on ligand-receptor interaction probability and B) leveraged inter-individual transcriptional networks to predict gene network changes with altered EGF expression in nuclei subpopulations. First, we computed CellChat on an individual sample basis to extract ligand receptor interaction probabilities for each individual which was then correlated to Rd/I to infer which cell-cell communication networks are affected with SkM insulin resistance (**Figure 4A, Supplementary Figure 4A).** EGF+ myonuclei ligand-receptor interactions that correlated with Rd/I were primarily targeted to FAP and immune cells (**Figure 4A**). EGF is the primary ligand for EGFR and in this data-set, EGF/EGFR ligand receptor interaction probability between EGF+ myonuclei and immune cells showed a negative correlation with Rd/I (**Figure 4B**). Meaning, increased interaction with EGF+ myonuclei and immune cells through EGF/EGFR signaling correlates with lower SkM insulin sensitivity. As there are multiple types of immune cells present in SkM^48^, we subclustered the immune cells into two broad classifications: Lymphoid and Myeloid cells (**Supplementary Figure 4B)** that were annotated with known marker genes (*IL7R, CD3E* & *BCL11B –* Lymphoid, *MRC1, F13A1, PDGFC-* Myeloid) (**Supplementary Figure 4C).** When CellChat was recomputed with this enhanced immune cell resolution it was found that Myeloid cells rather than Lymphoid cells had EGF/EGFR ligand-receptor interaction probability from EGF+ myonuclei that negatively correlated with Rd/I (**Supplementary Figure 4D).**

**Figure 4.**
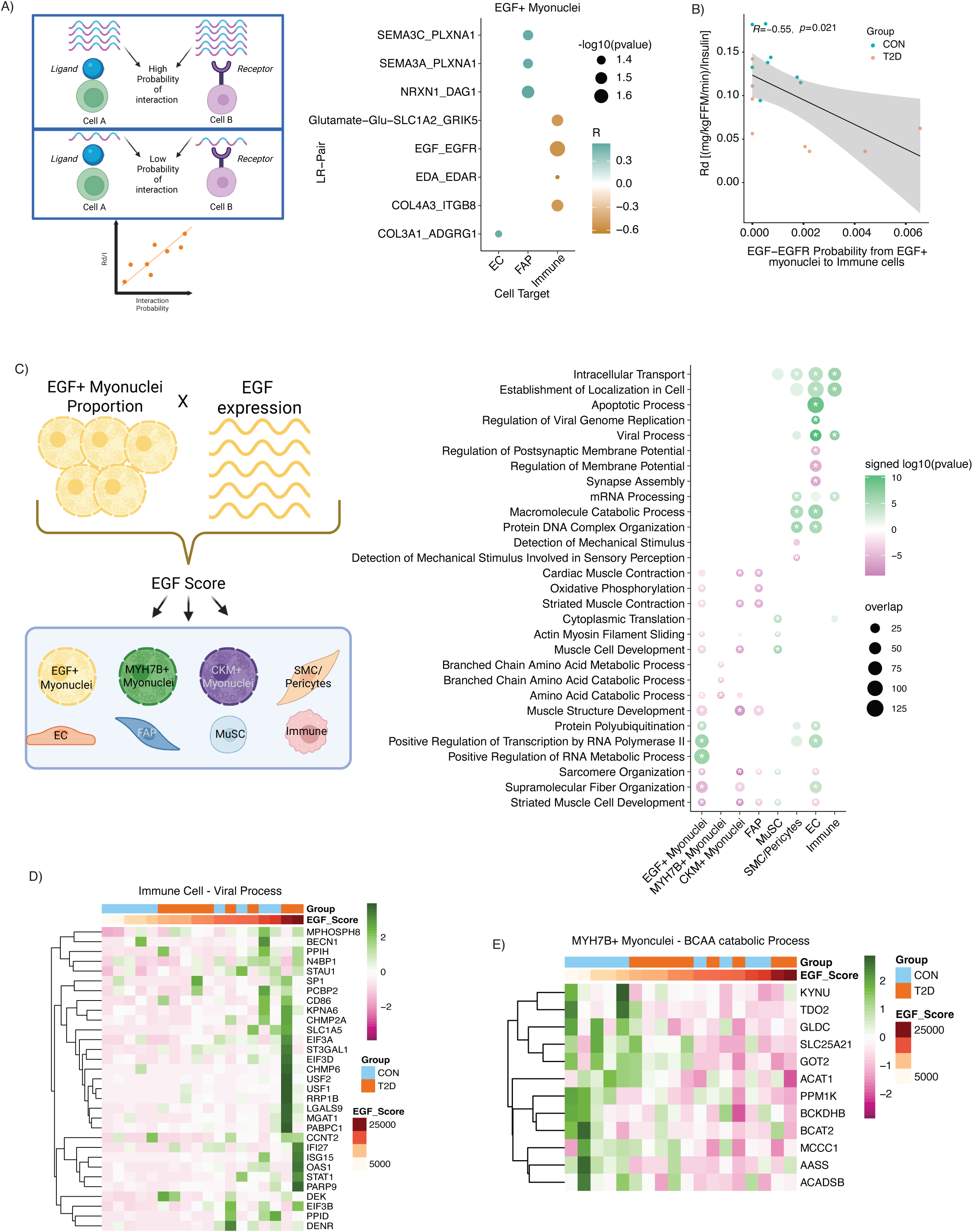
EGF+ Myonuclei cellular cross-talk. Association between ligand-receptor pair probability interaction and Rd/I for source nuclei type EGF+ Myonuclei (A). EGF/EGFR ligand receptor probability from EGF+ myonuclei to immune cells correlates negatively to Rd/I (B). The association between EGF score and transcriptional profiles of nuclei sub-types (C). EGF score positively correlates with viral process genes in immune cells (D). EGF score negatively correlates with BCAA catabolic process genes in MYH7B+ myonuclei (E).

Total EGF expression in SkM tissue is based on the proportion of EGF+ myonuclei and the overall expression of EGF in EGF+ myonuclei and therefore these two values were combined to generate an EGF score for each individual. This EGF score was then correlated to transcriptional profiles of nuclei sub-types to infer how the EGF ligand alters expression and function of cells with SkM (**Figure 4C)**. The number of correlating genes to each nuclei sub-type is shown in **Supplementary Figure 4E** and the full list of genes are included in **Supplementary Table 7.** Top pathways upregulated and downregulated with EGF score (**Supplementary Table 8**) are highlighted in **Figure 4C** and confirm known effects of EGF signaling. For example, EGF is known to activate skeletal muscle satellite cells and promote myoblast proliferation^49^, but EGF/EGFR signaling needs to be downregulated to stimulate myogenic differentiation^50^. Accordingly, these pathways are up and downregulated, respectively, in MuSC and EGF+/CKM+ myonuclei with EGF score (**Figure 4C**). In the immune cells genes related to viral processes were upregulated with a higher EGF score (**Figure 4D)** which is reflective of enhanced regulation of proliferation and activation of the cells. Within this gene-set were genes known to control T cell regulation (*PCBP2, CD86, LGALS9 & ISG15*)^51–54^ indicative of a more pro-inflammatory response. When these analyses were recomputed with higher immune cell resolution (Lymphoid and Myeloid) (**Supplementary Table 8**) viral process was not upregulated in either higher resolution immune cell type, (**Supplementary Table 8**) indicating the viral process gene signature is a combined response from all immune cells. Intriguingly, viral process was also upregulated in EC with higher EGF score; however, it was composed of different gene signatures to the immune cell viral process (**Supplementary Figure 4F**).

Interestingly amino acid and specifically branch chain amino acid (BCAA) catabolism gene signatures were negatively correlated with EGF score in MYH7B+ myonuclei only (**Figure 4E**). Elevated plasma BCAA is a common molecular signature observed with insulin resistance and T2D^55–57^ and is believed to be a consequence of impaired BCAA catabolism in SkM^58^. BCAA catabolism requires initial reversible transamination by BCAA aminotransferase (*BCAT1 & BCAT2*) followed by oxidative decarboxylation by the branched-chain alpha keto acid dehydrogenase complex (*BCKDHA & BCKDHB*). This complex is inactivated by BCKD kinase (*BCKDK*) and activated by Branched-Chain Alpha-Ketoacid Dehydrogenase Phosphatase (*PPM1K*). Downstream CoA compounds are further metabolized to acetyl-CoA and succinyl-CoA and incorporated into the TCA cycle^56,58^. Important genes involved in the initial BCAA catabolism steps (*BCAT2, BCKDHB & PPM1K*) and downstream CoA metabolizing steps (*MMCC1, ACAT1 & ACADSB*) are downregulated with higher EGF score in MYH7B+ myonuclei only (**Figure 4E**). Similar downregulation of BCAA catabolism genes in SkM is also observed in individuals with IR^12,59^.

## Discussion

SkM is a major site for post-prandial glucose disposal owing to its large tissue volume and the substantial glucose storage and oxidative capacity of myofibers. Interspersed among myofibers is a diverse collection of cells that can either assist or impair myofiber glucose uptake from the circulation^14^. Here, by integrating full-length single-nucleus transcriptional profiling of human SkM with gold standard metabolic phenotyping, we reveal SkM insulin resistance is associated with (i) the proportions of previously unrecognized myonuclear populations, ii) nucleus-specific molecular signatures from myogenic and non-myogenic cells and iii) intercellular signaling pathways between resident cell-types.

Skeletal muscle fibers have traditionally been classified as slow-twitch Type I fibers that have enhanced oxidative capacity and fast-twitch Type II fibers that are more glycolytic, with the latter being further divided into aerobic (Type IIa) and anaerobic (Type IIx) subtypes^60–62^. Until recently, fibers were classified on these sub-types based on the expression of singular myosin heavy chain markers (MYH1/MYH2/MYH7). However, pioneering work that combined advanced transcriptomics and proteomics workflows to individual myofibers revealed that fiber-type heterogeneity extends beyond traditional myosin heavy chain expression/abundance with metabolic, ribosomal and cell junction molecular features driving greater variation^37^. As myofibers are syncytia, their expression profiles are accumulative of all the myonuclei resident in the fiber. By performing snRNA-seq we extend transcriptional diversity beyond traditional myosin heavy chain to the myonuclei and highlight novel myonuclei populations; EGF+ myonuclei, MYH7B+ myonuclei and CKM+ myonuclei which were further validated with RNAScope. Previous snRNA-seq SkM research has clustered by traditional fiber typing but is constrained by gene body coverage capabilities from the restricted 3’ only technology^24,25,27–29,63^. We propose the superior gene detection capabilities generated from the 3’ and 5’ amplification and profiling used in this study resulted in novel myonuclei populations being identified. Interestingly CKM+ myonuclei were defined by an upregulation of mitochondrial and cytoplasmic translation genes which were primary transcriptional drivers defining fiber-type diversity in the work by Morena-Justicia et al.,^37^. In addition to identifying and validating distinct myonuclear populations, this work highlights their proportions relate to SkM insulin sensitivity in two independent datasets with MYH7B+ myonuclei being more metabolically favorable than EGF+ myonuclei.

Consistent with prior studies, comparisons between discrete control and T2D groups revealed limited and poorly coordinated molecular differences^43^, prompting our analyses to model transcriptional variation against continuous measures of insulin sensitivity. HOMA-IR is a commonly used insulin sensitivity index that is computed from fasting insulin and glucose and therefore is more reflective of hepatic rather than SkM insulin resistance^64^, even when using the non-linear HOMA2 model. A HE clamp quantifies whole body insulin sensitivity from glucose infusion rate (a.k.a M-value) which at high insulin doses is more reflective of peripheral glucose uptake rather than suppression of hepatic glucose production. The addition of a stable isotope-labelled glucose tracer during the HE clamp further allows for the direct quantification of peripheral glucose disposal (Rd) at physiological insulin doses and is therefore the gold-standard method of assessing SkM insulin sensitivity. Modelling transcriptional profiles against Rd/I uncovered nucleus-specific molecular signatures associated with SkM insulin resistance, with minimal overlap of genes and pathways across nucleus types. Notably, this nucleus-specificity was not observed when modelling to HOMA2-IR, and gene expression profiles associated with insulin sensitivity differed markedly between Rd/I and HOMA2-IR-based analyses. Together, these findings underscore the importance of gold-standard metabolic phenotyping and reveal distinct, nuclei-specific molecular features of SkM insulin resistance that would be overlooked by bulk transcriptomic approaches.

We identified *SLC39A14* encoding zinc transporter ZIP14, as a molecular feature associated with SkM insulin resistance in both CKM+ and EGF+ myonuclei. Whereas ZIP14 reduces glucose uptake by promoting insulin receptor degradation in hepatocytes^45^, we observed the opposite effect in SkM, where increased ZIP14 resulted in greater insulin-stimulated glucose uptake. ZIP14 KO mice have increased NF-κB in SkM^46^ likely due to impaired zinc-mediated suppression of inflammatory pathways. As IKK-β both activates NF-κB and impairs insulin signaling via serine phosphorylation of IRS-1^65^, it is plausible that ZIP14 mediated zinc transport downregulates NF-κB signaling, restores tyrosine phosphorylation of IRS-1 and therefore preserves insulin sensitivity. Intriguingly, ZIP14 expression is upregulated with inflammation^46^, and we observed ZIP14 strongest effects on insulin stimulated glycogen synthesis under lipid-induced insulin-resistant conditions, suggesting a protective role in inflammatory induced insulin resistance. In support of this interpretation, zinc supplementation improves glycemic control in individuals with T2D^66^; whether these benefits can be enhanced through ZIP14 modulation now warrants further investigation.

Impaired BCAA catabolism is an established signature of SkM IR and T2D^55–58^. Using a systems biology approach, we provide evidence that EGF signaling contributes to this metabolism signature, paralleling observations in colorectal cancer^67^. EGF/EGFR signaling increases in aging SkM^68^ and promotes senescence by secreting inflammatory cytokines via Ras-dependent pathways^69^. Improving BCAA catabolism through inhibition of BCKD kinase increases BCAA oxidation and reduces plasma BCAA levels^58^ and in some instances improves peripheral insulin sensitivity in mice and humans^70,71^. Together, these findings suggest that EGF-mediated suppression of BCAA catabolism may represent a previously underappreciated mechanism linking aging, inflammation and SkM insulin resistance and supports therapies that target EGF receptor inhibition. Consistent with this interpretation, EGFR-targeting tyrosine kinase inhibitors have recently been identified as high-priority candidates for pharmacological repurposing to improve insulin sensitivity^13^.

Despite leveraging advanced single-nucleus transcriptomics with gold-standard phenotyping of SkM insulin resistance to generate the first transcriptomics atlas of human SkM insulin resistance, several limitations should be acknowledged. To assess the transcriptional aberrations occurring with SkM insulin resistance we limited the effects of confounding factors by selecting participants with similar elevated BMI and age and reduced aerobic capacity. This design likely precludes detection of earlier transcriptional aberrations associated with obesity and aging and therefore future studies should incorporate lean and younger cohorts to capture these initial molecular alterations. Additionally, our sample size was modest and predominantly female. By deconvoluting larger and more diverse bulk RNA-seq datasets, we validated insulin resistance associated differences in myonuclear proportions and assessed age-, sex- and BMI effects. Single-nucleus transcriptomic profiles of SkM are dominated by myonuclei due to their abundance within myofibers, thus limiting resolution of mononuclear cell populations. While single-cell RNA-seq allows improved resolution of these population, it is at the expense of myonuclear representation. Emerging spatial transcriptomics approaches have the potential to profile all cell-types within SkM while preserving their spatial relationship to neighboring cells.

In conclusion, these findings establish myonuclear diversity as a key determinant of SkM insulin sensitivity and highlight the importance of coupling high-resolution transcriptomics with precise metabolic phenotyping to uncover nucleus-specific dysregulation that is obscured with bulk analyses. This nucleus-centric view of SkM insulin resistance drives new avenues for targeted metabolic interventions to prevent or delay T2D.

## Methods

### Participant characteristics

Samples for this study were collected as part of a larger clinical study investigating the effects of weight loss and exercise on insulin sensitivity, body composition, aerobic capacity and muscle strength in older individuals with obesity^72^. Only baseline samples were used for this manuscript. All participants were weight stable over the last 6 months, physically inactive (≤1 continuous exercise session/week) and non-smokers. Exclusion criteria included clinically significant cardiovascular disease including history of myocardial infarction within the past year; peripheral vascular disease; hepatic, renal, muscular/neuromuscular, or active hematologic/oncologic disease; the presence of bruits in the lower extremities; history of pulmonary emboli; peripheral neuropathy; anemia; and substance abuse. Participants on the following medications were excluded anticoagulants, glucocorticoids, thiazolidinediones, or insulin. Full list of medications can be found in Supplementary Table 9. The study was approved by AdventHealth Institutional Review Board and carried out in accordance with the Declaration of Helsinki. Participants provided written informed consent to partake in the study. All samples were taken following an overnight fast.

Samples were selected of individuals with and without T2D with a range in SkM insulin sensitivity to determine the effects of SkM insulin resistance and T2D on nuclei specific SkM transcriptome. Full phenotypic testing of participants including aerobic capacity (VO_2peak_ test), body composition (DEXA) and SkM insulin sensitivity (hyperinsulinemic euglycemic (HE) clamp) have been described in detail previously^72^. The non-linear HOMA2-IR variable was calculated using the HOMA2 model (www.OCDEM.ox.ac.uk). As SkM insulin sensitivity is one of the key phenotypical traits of these analyses, it is also explained in detail below.

#### Hyperinsulinemic-Euglycemic Clamp

Participants arrived at the Translational Research Institute the evening prior to the HE clamp and consumed a standard meal before staying overnight in the metabolic ward. Following the overnight fast, an intravenous catheter was placed in the antecubital vein for subsequent insulin, glucose, and stable isotope infusions. A primed infusion of [6,6-2H2] glucose was ran constantly throughout the HE clamp. An additional catheter was placed on the contralateral arm on a heated hand vein to attain arterialized blood samples for blood glucose determination and for [6,6-2H2] glucose enrichment during the insulin and glucose infusions. After a 2.5-hour baseline period, insulin was infused at 40 mU/m^2^ min and continued for 4 hours. Glucose was measured at 5-minute intervals and maintained at 90 mg/dL. A 2mL blood sample was collected at 0, 30, 60, 100, 110, and 120 minutes and every 10 minutes during the last 30 minutes of the HE clamp for determination of [6,6-2H2] glucose enrichment using gas chromatography–mass spectrometry. Insulin samples were also drawn at multiple time points. Rates of glucose disposal (R_d_) were calculated by nonsteady-state equations based on plasma [6,6-2H2] glucose enrichment^73,74^. SkM insulin sensitivity (R_d_/Insulin) was assessed as the rate of glucose disposal (mg/kgFFM/min) accounting for plasma insulin during steady state. Indirect calorimetry using a metabolic hood (ParvoMedics TrueOne 2400 system, US) was performed with urinary nitrogen corrections to calculate rates of fasting and insulin-stimulated glucose oxidation and NOGD.

### Skeletal Muscle Biopsy

Following an overnight fast and prior to the HE clamp a percutaneous muscle biopsy of the *vastus lateralis* was performed using previously published methods^75^. A biopsy sample was taken 10–15 cm above the knee under local anesthesia with a 5-mm Bergstrom^76^ needle and suction. Following removal of excess blood, fat and connective tissue, a portion was snap-frozen in liquid nitrogen for downstream analyses including snRNA-Seq. Another portion of the tissue was mounted in Tissue-Tek (Sakura Finitek, US) and frozen in liquid nitrogen cooled isopentane for immunohistochemistry analyses.

#### Single nuclei RNA-Seq

Nuclei was isolated from frozen SkM (∼10mg) using a modified approach to our previously published protocols^77,78^. Briefly SkM was pulverized under liquid nitrogen prior to being homogenized in 1mL of homogenization buffer (5mM MgCl2, 25mM Tris Buffer pH 8.0, 25mM KCL, 250mM sucrose, 1μm DDT, 1 x protease inhibitor, 0.2 U/μL SUPERase · In RNase Inhibitor (Thermofisher Scientific) in nuclease-free water) with a glasscol homogenizer. After addition of Triton-X100 (0.1% v/v) the homogenate was incubated on ice with regular vortexing. Samples were then filtered through a 100μm and 40μm strainer (BD Falcon), centrifuged at 1000g for 10 min at 4°C and then re-suspended in 500μL nuclei isolation medium (5mM MgCl2, 25mM Tris Buffer pH 8.0, 25mM KCL, 1 mM EDTA, 0.2 U/μL Ribolock RNAase inhibitor, 1% BSA in nuclease-free water) before being centrifuged at 1000g for 10 min at 4°C. The sample was re-suspended in 500μL of 1%BSA-PBS (-/-) with 0.2 U/μL Ribolock RNAase inhibitor and filtered through a 30μm strainer (Miltenyi Biotec). Nuclei was stained with Hoechst 33342 (ReadyProbes Cell Viability Imaging Kit, Thermofisher Scientific) for 15 min prior to counting with a countess II automated cell counter (Thermofisher Scientific).

Single-nuclei suspension (40K/mL) was aliquoted into 8 wells of a 384-well source plate (Takara Bio, USA, San Jose, CA) and dispensed using an iCELL8 MultiSample NanoDispenser (Takara Bio, USA) onto an iCELL8® 350v Chip (Takara Bio, USA). After dispense, the nanowells of the chip were imaged using the iCELL8 Imaging Station to identify nanowells containing a single nucleus that were selected for downstream library generation. Additionally, 50 nanowells that did not contain any nuclei were also selected for downstream library generation to serve as an ambient control. After imaging, the chip was subjected to freeze-thaw to lyse the nuclei and then selected nanowells were subjected to first-strand cDNA synthesis initiated by oligo dT primer (SMART-Seq Pro CDS), followed by template switching with template switching oligo (SMART-Seq Pro oligonucleotide) for 2^nd^ strand cDNA synthesis, before unbiased amplification of full-length cDNA. Bead-Linked Transposome (Illumina, San Diego, USA) was used to tagment full-length cDNA before amplification with forward (i5) and reverse (i7) indexing primers. Each single nucleus was indexed by a unique combination of 1 of 72 forward and 1 of 72 reverse indexing primers allowing for downstream identification. Collected cDNA was purified twice using a 1:1 proportion of NucleoMag NGS Clean-Up and Size Select beads (Takara Bio, USA). cDNA was further amplified according to manufacturer’s instructions and purified again at a 1:1 proportion of NucleoMag beads. The resultant cDNA library was assessed for concentration by fluorometer (Qubit, Thermofisher Scientific) and quality by electrophoresis (Agilent Bioanalyzer high sensitivity DNA chips). Libraries were sequenced in pairs with NovaSeq X Plus PE150 targeting 375G of data per lane which downstream equated to an average 415,484 barcoded reads per nuclei.

#### Fluorescent in situ hybridization (FISH)

Fluorescent in situ hybridization (FISH) was performed with RNAscope multiplex fluorescent kit v2 (ACDbio, CA, USA) and a 4-plex ancillary kit (ACDbio) with probes against MYH7B (456611- C1), EGF (605771-C2), MYH2 (504731-C3) and MYH7 (508201-C4) according to the manufacturer’s protocol. SkM samples were sectioned at 10 µm at -20°C onto Superfrost^TM^ Plus slides (Fisherbrand, USA) and dried at -20°C for 60-120 min prior to storage at -80°C. Sections were fixed in 4% parafomaldahyde, washed twice in 1x PBS and dehydrated with sequential incremental ethanol incubations. After the slides had dried they were incubated in hydrogen peroxide (ACDbio) for 10 min at room temperature, washed twice in distilled water and incubated with protease IV (ACDbio) for 30 min at room temperature, before being washed again in distilled water. Probes were combined into one solution and hybridized on the slides at 40°C for two hours. To assess tissue quality a positive 4-plex control probe (#321801) and 4-plex negative control probe (#321831) was performed simultaneously. Sequential hybridize amplification were performed before each HRP probe signal was sequentially developed. Fluorophore; 570 TSA vivid labelled the MYH7B-C1 probe, 650 TSA vivid labelled the EGF-C2 probe, 520 TSA vivid labelled the MYH2-C3 probe and Opal Polaris 780 (Akoya) labelled the MYH7-C4 probe. After signal development, sections were stained with wheat germ agglutin (WGA)- 680 for 30 min to aid with visualization and then stained DAPI for 5 min to detect nuclei. Slide were flicked dry before a cover slip was mounted with ProLong Gold Antifade Mountant (Thermofisher). Images were obtained at 40x magnification on a Sp5 confocal microscope (Leica, Germany). Samples from 5 individuals from the same cohort as the snRNA-seq analyses were used with an average (SD) of 7 ± 1 40x images analyzed per sample. This resulted in an average (SD) 462 ± 57 nuclei analyzed per sample.

Images were analyzed using QuPath-0.5.1x64. Nuclei were detected with the cell detection function and intensity measures for each channel within the nuclei were extracted for downstream analyses. A percentage expression score for each nuclei was calculated for all markers used (MYH2, MYH7, MYH7B and EGF). Nuclei with a percentage score > 50% for a marker was classified as that that marker e.g. MYH2+. Nuclei with percentage score >40% for two markers were classified as those two markers e.g. MYH7B+/MYH7+. Nuclei with percentage score >30% for 3 markers were classified as those 3 markers e.g. MYH7+/MYH2+/EGF+. The proportion of different nuclei populations was calculated for each sample across all images.

### Bioinformatic analyses

Demultiplexing, mapping, alignment and counting of the single nuclei RNA-Seq libraries were performed using CogentAP™ Analysis Pipeline (Takara Bio, USA). GRCh38 was used as the genome reference and included 58735 transcripts. Nuclei were initially removed if they had; < 500 genes, >15,000 genes, < 10,000 counts, > 30% mitochondrial reads, >15% protein coding mitochondrial reads, > 30% rRNA reads or nuclei complexity < 0.6. To remove low expressed genes from clustering analyses, genes with an average count less than 0.1 were removed. Data was normalized in Seurat with SCTransform^79,80^. Initially each sample was clustered separately using 3000 highly variable genes to detect sample dependent cell clusters. Data was then adjusted for ambient RNA with decontx^81^ using cell clusters labels as the Z parameter and using ambient wells as the background reference. We next compared multiple integration methods; unadjusted, STACAS, fastMNN, Seurat (RPCA) and harmony (Andreatta & Carmona, 2021; Hafemeister & Satija, 2019; Hao et al., 2021; Korsunsky et al., 2019; Zhang et al., 2019). For each integration 5000 highly variable genes were used and where applicable the number of PCA dimensions used for integration was 1:20 and the number of PCA dimensions used for UMAP projection and clustering was (1:10). Adjusted rand index (ARI)^85^ was calculated using the *adj.rand.index* function in *pdfCluster* v1.04. by providing a list of participant IDs and identified cell clusters. An ARI closer to 0 indicates random clustering of samples among the different cell types and good batch correction. Average Local Inverse Simpson’s index (LISI) scores were estimated using the *compute_lisi* function in *lisi* v1.0^83^ by providing a list of the UMAP embeddings along with the sample ID and cell clusters to estimate how well-mixed donors are across the cell clusters. As there were 18 samples, a score closer to 20 would indicate the cells are entirely mixed among participants whereas 0 would indicate the cells are not mixed well. Harmony had the highest LISI score and lowest ARI score indicating the samples were well integrated (**Supplementary Table 10**) and this was used for downstream analyses. The LISI score for harmony was 8.6 indicating although it was the highest and well integrated batch correction method there still biological variability among the samples.

To ensure erroneous conclusions were not made from reminiscent ambient profiles after decontX adjustment, ambient gene culprits were determined by having average count expression > 100 and being present in at least 50% of the ambient control nanowells (**Supplementary Table 11**) which resulted in 399 ambient genes. These cutoffs were determined by histogram plots showing the majority of ambient RNAs are highly expressed and are in the majority of nanowells. 163 of the genes (35%) were cluster specific (upregulated in a cluster) with the largest origin coming from the myonuclei. This ambient gene list was used as a QC for interpreting downstream results.

To identify nuclei types markers differential gene expression analyses for nuclei clusters was performed using a Wilcoxon rank sum test with Seurat’s “FindMarkers” function with a FDR cut off of < 0.05, log_2_ FC > 0.25 or < -0.25 and expressed in >25% of nuclei in that cluster.

#### Myonuclei heterogeneity

Myonuclei fiber-typing on CKM+ myonuclei was calculated based on the expression of traditional myosin heavy chains (MYH1, 2, 7) similar to previous analyses^27^. For each myonuclei the percentage expression of each myosin heavy chain was calculated. An additional score was calculated based on the combined expression of MYH2 and MYH1 representing combined Type II nuclei. Type I myonuclei were assigned if the ratio of MYH7 to MYH2/1 expression exceeded more than 1 and if the MYH2 score was less than 1. If the ratio of MYH7 to MYH2/1 exceeded 1 and the MYH2 score was greater than 1, myonuclei were classified as hybrid Type I/IIA. Afterwards, myonuclei were classified as Type IIa if the ratio between MYH2 to MYH1 was greater than 1.25 and as Type IIX if the ratio was less than 0.75. If the ratio between MYH2 to MYH1 was between 0.75 and 1.25 myonuclei were classified as hybrid Type IIA/IIX.

To compare transcriptional profiles between myonuclei populations Seurat FindMarkers() function was run comparing each myonuclei population to each other. Upregulated genes (adjusted p value < 0.05 & logfc > 1) for each myonuclei population was used as input pathway analysis. Pathway analysis was conducted on gene ontology: biological process genesets^86,87^ using TMsig (V1.0.0)^88^ and hypeR (V2.4.0)^89^. TMsig filters pathways for background gene expression and then reduces redundancy by filtering pathways with similar genesets. A hypergeomtric test was then conducted on filtered gene-set and upregulated genes in the HypeR package.

#### Statistics

Pearsons correlation between phenotypical traits and nuclei proportions were conducted with R Package Hmisc (V5.2-2) rcorr() function. For comparisons of phenotypical data and nuclei proportions between the CON and T2D, an unpaired *t*-test was used to detect differences in normally distributed data, and Mann–Whitney *U* test was used for non-normally distributed data using Rstatix R package (V0.7.2)

#### Deconvolution of publicly available bulk RNA-Seq datasets

A cell signatures matrix was created by aggregating counts across cell types and filtering to 5000 HVG that was used for integration and clustering. Our previous analyses concluded algorithm dtangle to be the most accurate method for deconvoluting bulk RNA-seq using full-length snRNA-seq data^90^. Bulk RNA-Seq data sets were converted to counts per million filtered for genes that had expression in at least 50% of samples and then deconvoluted using deconvolute() function in granulator with method dtangle. The output provides estimated proportions of each nuclei type which was used to calculate the ratio of MYH7B+ myonuclei to EGF+ myonuclei for downstream comparisons.

##### Datasets

*CALERIE*- This study was conducted on adults (21-50 years) without obesity and investigated the effects of 25% calorie restriction (CR) compared to ad libitum control over a 2 year period^91^. Deconvolution was performed on RNA-seq data from SkM biopsies of the *vastus lateralis* at baseline and 24 months which included 88 baseline samples (n = 55, CR) and 30 24-month samples (n = 20, CR)^92^.

*Trim et al., 2022*- This is a cross-sectional study comparing older (60-85 years) males (n =1 2) to younger (20-35 years) males (n =12) with similar levels of leanness and activity levels^40^. RNA-seq was performed on SkM biopsies from the *vastus lateralis*.

*Mahmassani et al., 2019*- This study compared the effects of 5-day bed rest on older (60-75 years, n = 19) and younger (18-35 years, n = 9) healthy individuals^41^. RNA-Seq was performed on SkM biopsies from the *vastus lateralis* pre and post intervention.

*GTEx-* The adult Genotype Tissue Expression (GTEx) study obtained RNA-seq data from the *gastrocnemius* of 818 decreased individuals^93^. The number of samples per age group and sex is displayed in the Table 2.

**Table 2:** Number of SkM samples analyzed from GTEx study.

| Age-Group (years) | Female | Male |
| --- | --- | --- |
| 20-29 | 23 | 48 |
| 30-39 | 18 | 46 |
| 40-49 | 50 | 71 |
| 50-59 | 85 | 176 |
| 60-69 | 82 | 191 |
| 70-79 | 8 | 20 |

*Devarshi et al., 2022*- This study investigated the effects of BMI on acute exercise responses in vastus lateralis using RNA-seq^42^. For deconvolution analyses we only used the baseline samples to compare lean individuals (n = 14) to individuals with overweight/obesity (n = 15).

#### Pseudobulk Analyses

##### Differential gene expression analyses

Single cell data was converted to SingleCellExperiment (V1.28.1) for downstream pseudobulk differential expression analyses with DESeq2 (V1.46.0) For each sample and nuclei-type, a pseudobulk counts matrix was generated by aggregating raw counts. Genes were removed per nuclei-type pseudobulk matrix if all values were 0 or if values were less than 1 in at least half of the samples. Data was normalized with the DESeq() function and differential gene expression analyses was conducted between T2D and CON with FFM as covariate. Sex showed little impact on variance and was therefore not used as a covariate. Upset plots from R package ComplexHeatmap were generated to show overlapping DEGs upregulated and downregulated with T2D. Pathway analysis was conducted on gene ontology: biological process genesets^86,87^ using TMsig^88^ and hypeR^89^. Normalized pseudobulk counts from the DESeqDataSet without FFM as a covariate were saved for downstream analyses.

##### Correlation analyses

Kendall rank correlational analyses between nuclei specific gene expression profiles (normalized pseudobulk counts) and skeletal muscle insulin sensitivity (Rd/I) was conducted using the cor.test() function from the stats r package (V4.4.2). For genes with a significant correlation to Rd/I (non-adjusted *P* < 0.05), empirical p values were calculated by permuting the Rd/I labels a 1000 times. Pathway analysis was conducted on gene ontology: biological process genesets^86,87^ using TMsig^88^ and hypeR^89^.

##### Cell Chat analyses

Cell-cell communication analysis was performed using CellChat (V 2.2.0)^94^ using the CellChatDB.human database on each individual sample. Over expressed genes and interactions were identified with the identifyOverExpressedGenes() and identifyOverExpressedInteractions() functions. Cell-cell communication was inferred by assigning each interaction a probability value using the computeCommunProb function with the default ‘triMean’ option and using raw data rather than smoothed data. Communication was then filtered to have a minimum of 10 cells and cell-cell communication at a signaling pathway level was calculated with computeCommunProbPathway(). The communication probability was then summarized with aggregateNet(). Ligand receptor interactions were filtered to have at least 3 samples with non-zero probability values. Communication probability per ligand-receptor interaction for each sample was then correlated to Rd/I and filtered for significance (*P* < 0.05).

### Human SkM cell culture

Isolation of primary human satellite cells, differentiation into myotubes and measures of glucose oxidation and glycogen synthesis rates as basal levels and following insulin stimulation with and without palmitate induction have recently been described in detail^47^. RNA was extracted from the myotubes with RNeasy fibrous tissue mini kit (Qiagen) and poly(A) libraries were generated and sequenced on Illumina 10plus. Counts were normalized to CPM using DESeq2. Pearson correlation was used to assess the relationship between *SLC39A14* expression and glucose uptake indices.

### Mouse SkM cell culture

#### Mouse Myoblast harvest

All animal procedures were approved by the University of Florida Institutional Animal Care and Use Committee and followed guidelines of the National Institutes of Health. ZIP14 KO and WT (C57BL/6) were bred at the University of Florida animal care services facility as previously described^46,95,96^ and were approximately 11 weeks old at the time of euthanasia. Mice were anesthetized with isoflurane and humanely euthanized by exsanguination via cardiac puncture. Quadricep and gastrocnemius SkM was harvested and minced in plating media (25% low glucose DMEM (with antibiotics, Gibco, US)), 25% Ham’s F-12 (with antibiotics, Invitrogen, US), 40% Heat-Inactivated FBS (Invitrogen, US), 10% Amniomax (Thermofisher, US). SkM fragments were then transferred into large droplets of plating media on Matrigel (Corning, US) and collagen (Gibco, US) pre-coated dishes and incubated for ∼ 7 days at 37°C, 5% CO_2_ for tissue attachment. Upon visible outgrowth myoblast cells were harvested in PBS and collected into growth media (25% low glucose DMEM (with antibiotics), 25% Ham’s F-12 (with antibiotics), 20% Heat-Inactivated FBS, 10% Amniomax). The plate was re-incubated with plating media, and the myoblast harvest was repeated after 2 days. Harvested myoblasts were centrifuged at 500g for 5 min, resuspended in growth media and plated for growth over 2 passages before cryopreservation.

#### Glucose Uptake in differentiated myotubes

Myoblasts were seeded at 200K in growth medium (25% low glucose DMEM (with antibiotics), 25% Ham’s F-12, 20% heat-inactivated FBS, 10% Amniomax) in 12 well-plates and at 100% confluency differentiated into myotubes over 5-7 days. Differentiation was confirmed visually and with Myh2 and Myh3 gene expression.

Following 3-4 hours of serum starvation, cells were washed twice with warm PBS and then incubated in triplicates for 30 min at 37°C and 5% CO_2_ with either; transport buffer (115mM NaCl, 2.6mM KCl, 1.2mM KH_2_PO_4_, 10mM NaHCO_3,_ 10mM HEPES, 0.1% w/v BSA), transport buffer with 100nM insulin or transport buffer with 20 µM Cytochalasin B to assess non-specific glucose incorporation. Label solution containing 0.44 µCi/mL ^3^H-Deoxyglucose (Perkin Elmer) and 20µm 2-Deoxy-D-glucose (Sigma Aldrich) was added to each well and incubated for 20-30min at 37°C and 5% CO_2._ Cells were washed twice with ice-cold PBS and lysed with 0.05 M NaOH. Cell lysates were added to scintillation fluid (Opti Flour) and ^3^H-Deoxyglucose levels were quantified with a scintillation counter. Protein concentration was measured in the remaining cell lysate using BCA protein assay (BioRad). Basal and insulin stimulated glucose uptake was corrected for non-specific glucose incorporation and adjusted to protein concentration. Due to the low sample size (n = 3, per group) the differentiation and glucose uptake experiments were repeated and data was combined for analyses. Different experimental batches are annotated in the figures.

#### Gene expression

RNA was extracted using RNAeasy Mini Kit (Qiagen, US) and real-time PCR was performed using a Viia7 sequence detection system (Life Technologies, US) and Taqman technology for relative gene expression quantification using the following parameters: one cycle at 50°C for 2 min, one cycle at 95°C for 10 min followed by 40 cycles of 95°C for 15 seconds and 60°C for 1 min. The primer/probes (integrated DNA technologies, US) used are shown in table 3.

**Table 3:** Primer Probes used in ZIP14 KO mice experiment.

| Gene | Assay ID | Exon Location |
| --- | --- | --- |
| *Zip14* | Mm.PT.58.28702454 | 6-8 |
| *Myod* | Mm.PT.58.8193525 | 1-3 |
| *Myh2* | Mm.PT.58.13701815 | 39-40 |
| *Myh3* | Mm.PT.58.32959978 | 1-3 |
| *Rplp0* | Mm.PT.58.43894205 | 5-6 |
| *Ppia* | Mm.PT.39a2.gs | 4-5 |

| Gene | Assay ID | Exon Location |
| --- | --- | --- |
| *Zip14* | Mm.PT.58.28702454 | 6-8 |
| *Myod1* | Mm.PT.58.8193525 | 1-3 |
| *Myh2* | Mm.PT.58.13701815 | 39-40 |
| *Myh3* | Mm.PT.58.32959978 | 1-3 |
| *Rplp0* | Mm.PT.58.43894205 | 5-6 |
| *Ppia* | Mm.PT.39a2.gs | 4-5 |

Relative gene expression was calculated by transforming ΔCt to the linear scale and the Livak method was used to assess changes in gene expression from pre to post myotube differentiation.

Glucose uptake was analyzed using a linear-mixed-effects model accounting for repeated measures (basal vs insulin) between genotypes, with sample as a random intercept and pairwise post-hoc Tukey comparisons. ZIP14 KO mice have impaired myogenesis^46^ and because myoblasts have enhanced basal glucose uptake^97^ the model was also adjusted for differentiation markers *Myh2* and *Myod1* expression. Preliminary testing revealed *Myh3* had no relationship with glucose uptake so was omitted as a covariate. Model-adjusted glucose uptake values were used for visualization.

### Data and Code Availability

The snRNA-Seq data generated during this study have been deposited to NCBI GEO and will be made available upon publication. Code Availability: Code used to process and analyze data in this paper have been deposited to GitHub and is publicly available at https://github.com/KWhytock13/skm-points as of the date of publication.

## Acknowledgements

This work was supported by the National Institute of Health, K99 DK135915 (KLW), R01 AG021961 (BHG), R01 DK120322 (LMS and JAH), DK 94244 (RJC) and Boston Family Endowment Funds of the University of Florida (RJC).

The authors thank the contributions of our study participants and acknowledge the expertise and assistance of the clinic and laboratory staff at Translational Research Institute- AdventHealth.

## Declaration of interests

MJW is considered a non-executive minority partner with Arch Venture Partners, Inc. The individual companies supported and funded by Arch have no decision-making function or stakeholder involvement.

## Author Contributions

Conceptualization – KLW; Data Curation – KLW, AD, JV, MH, MV, MAGM, CHR, FRJR, FM, PK, NTB, YS; Formal Analyses – KLW, AD; Funding Acquisition – KLW, RJC, JAH, LMS, BHG; Investigation- KLW, AD, JV, MH, MV, MAGM, CHR, FRJR, FM, PK, NTB, YS; Methodology- KLW, AD, JV, MH, PK, NTB, YS, MJW; Project Administration – KLW; Resources – RJC, MJW, JAH, LMS, BHG; Software- KLW, YS; Supervision- KLW, AD, MJW, RJC, JAH, LMS, BHG; Validation- KLW, AD; Visualization – KLW; Writing – Original Draft – KLW; Writing – review & editing – KLW, AD, JV, MH, MV, MAGM, CHR, FRJR, FM, PK, NTB, YS, MJW, RJC, JAH, LMS, BHG.

---

**Supplementary Figure 1.**
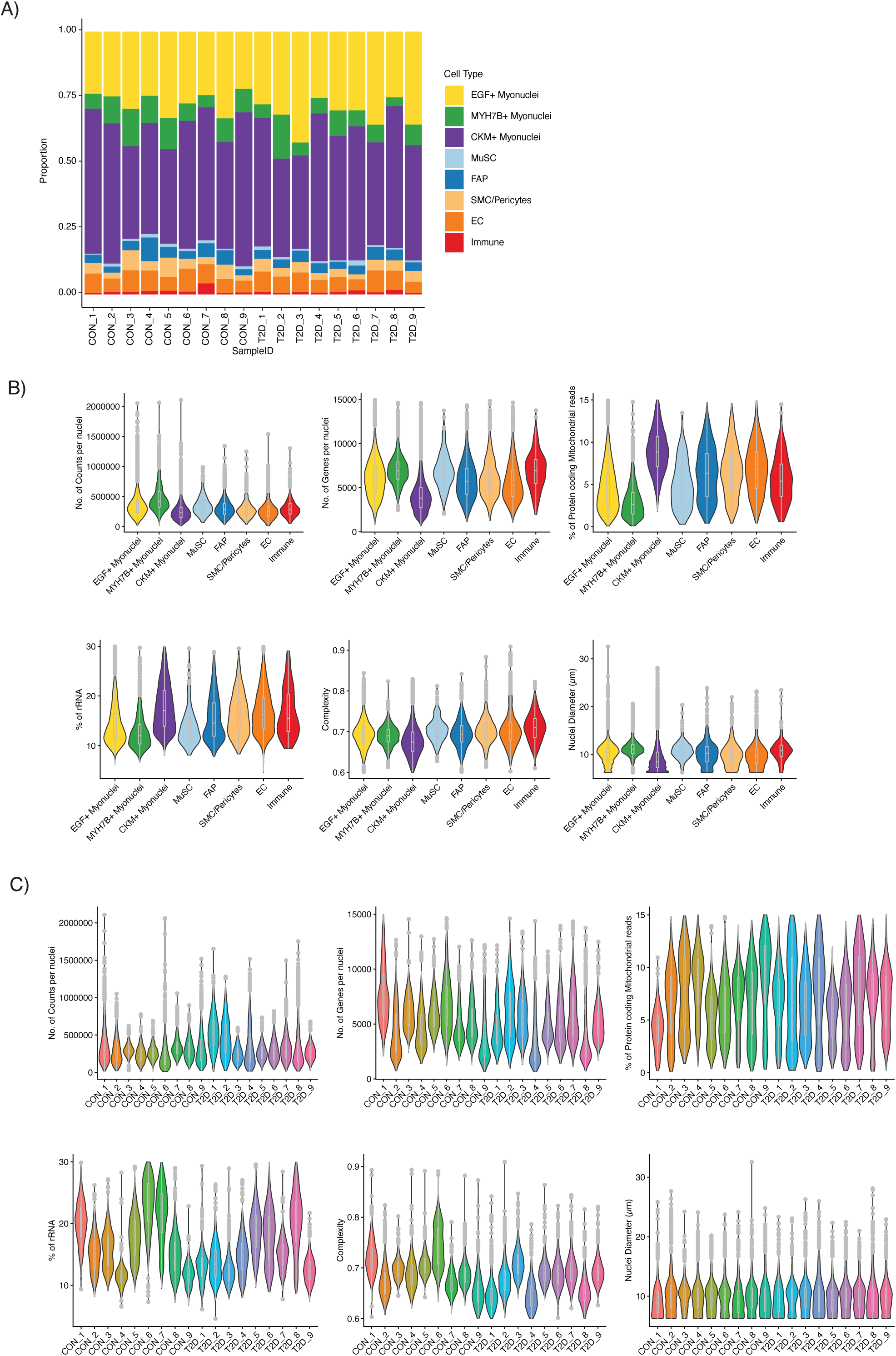
Supplement to Figure 1. Quality Control metrics for snRNA-Seq. Nuclei-type proportions for each individual sample highlights one cluster is not being driven by a single sample (A). Violin plots of quality control metrics show similar distribution among nuclei-types (B) and samples (C).

**Supplementary Figure 2.**
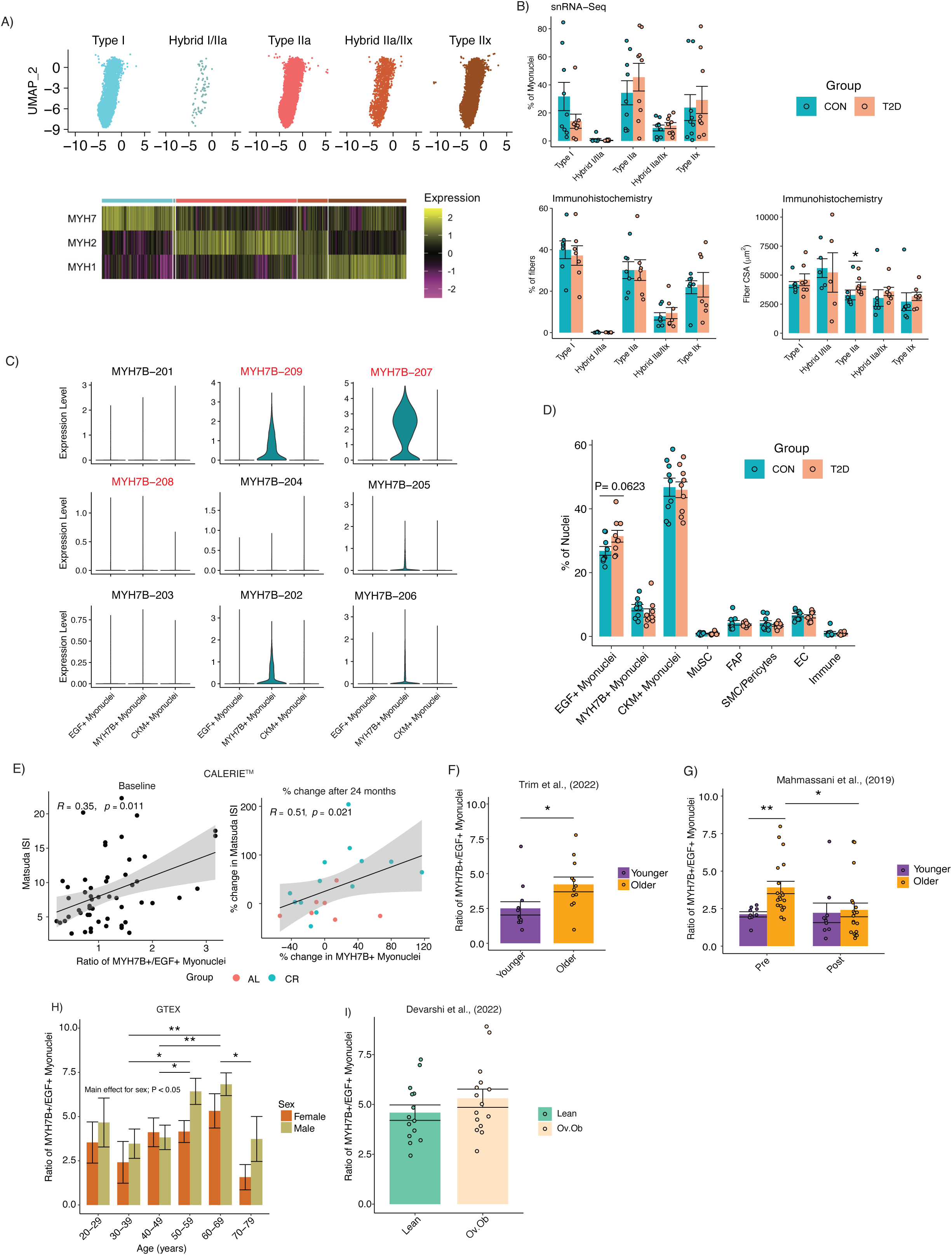
Supplement to Figure 2. CKM+ myonuclei can be characterized as Type I, hybrid I/IIa, Type IIa, Hybrid IIa/IIx and Type IIx when stratified based on traditional myosin heavy chain expression (A). Differences in traditional myosin heavy chain myonuclei/fiber proportions from snRNA-seq and immunohistochemistry analyses (B). MYH7B transcript expression among the different myonuclei populations. Transcripts in red indicate non-protein coding transcripts (C). Nuclei composition differences between T2D and Control (D). In the CALERIE^TM^ study baseline Matsuda ISI correlated positively with ratio of MYH7B+/EGF+ myonuclei ratio. After the 24-month intervention % change in Matsuda ISI positively correlated with % change in MYH7B+ Myonuclei (E). Older individuals had a greater ratio of MYH7B+/EGF+ Myonuclei compared to younger individuals in a study by Trim et al., (2022) (F). In a bed-rest study, Older compared to Younger individuals had a greater ratio of MYH7B+/EGF+ Myonuclei at baseline, which decreased to comparable levels to the younger group post-intervention in a study by Mahmassani et al., (2019) (G). In the GTEX study the ratio of MYH7B+/EGF+ Myonuclei increased with age up to 60-69 years before decreasing in the 70-79 age bracket with males having a higher MYH7B+/EGF+ Myonuclei ratio (H). The were no significant differences in the ratio of MYH7B+/EGF+ myonuclei between lean and individuals with overweight/obesity (Ov.Ob) (I). *, *P* < 0.05. **, *P* <0.05.

**Supplementary Figure 3.**
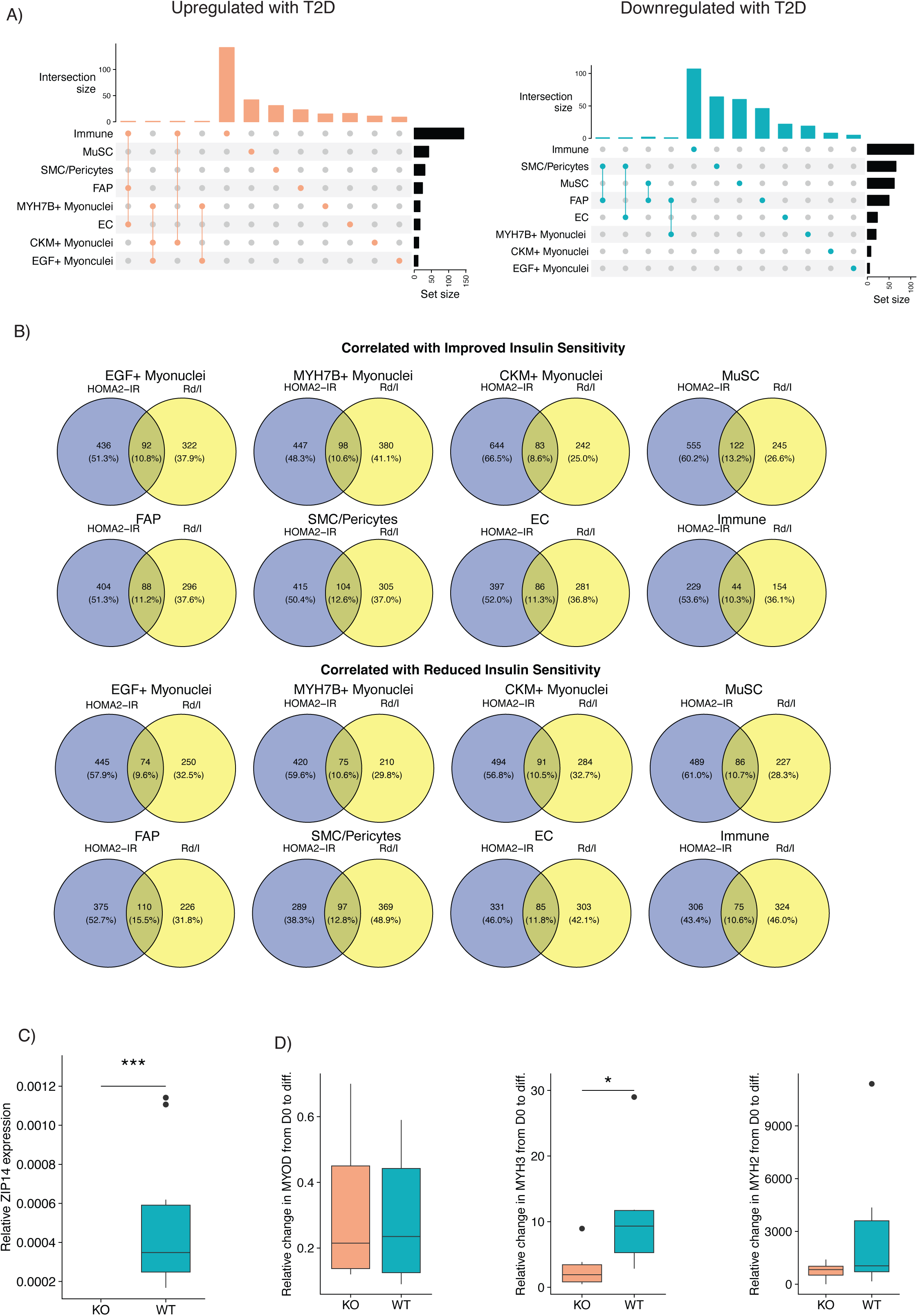
Supplement to Figure 3. Upset plots highlighting the number of differentially expressed genes upregulated and downregulated with T2D for each nuclei type (A). Venn diagrams of the overlap between genes correlated with improved insulin sensitivity (positive correlation to Rd/I and negative correlation to HOMA2-IR) and with genes that correlated with reduced insulin sensitivity (negative correlation to Rd/I and positive correlation to HOMA2-IR) among the different nuclei populations (B). Zip14 expression in SkM of WT and ZIP14 KO mice (D). Changes in myogenic markers during *in vitro* myotube differentiation of WT and ZIP14 KO muscle (D). *, *P* < 0.05.

**Supplementary Figure 4.**
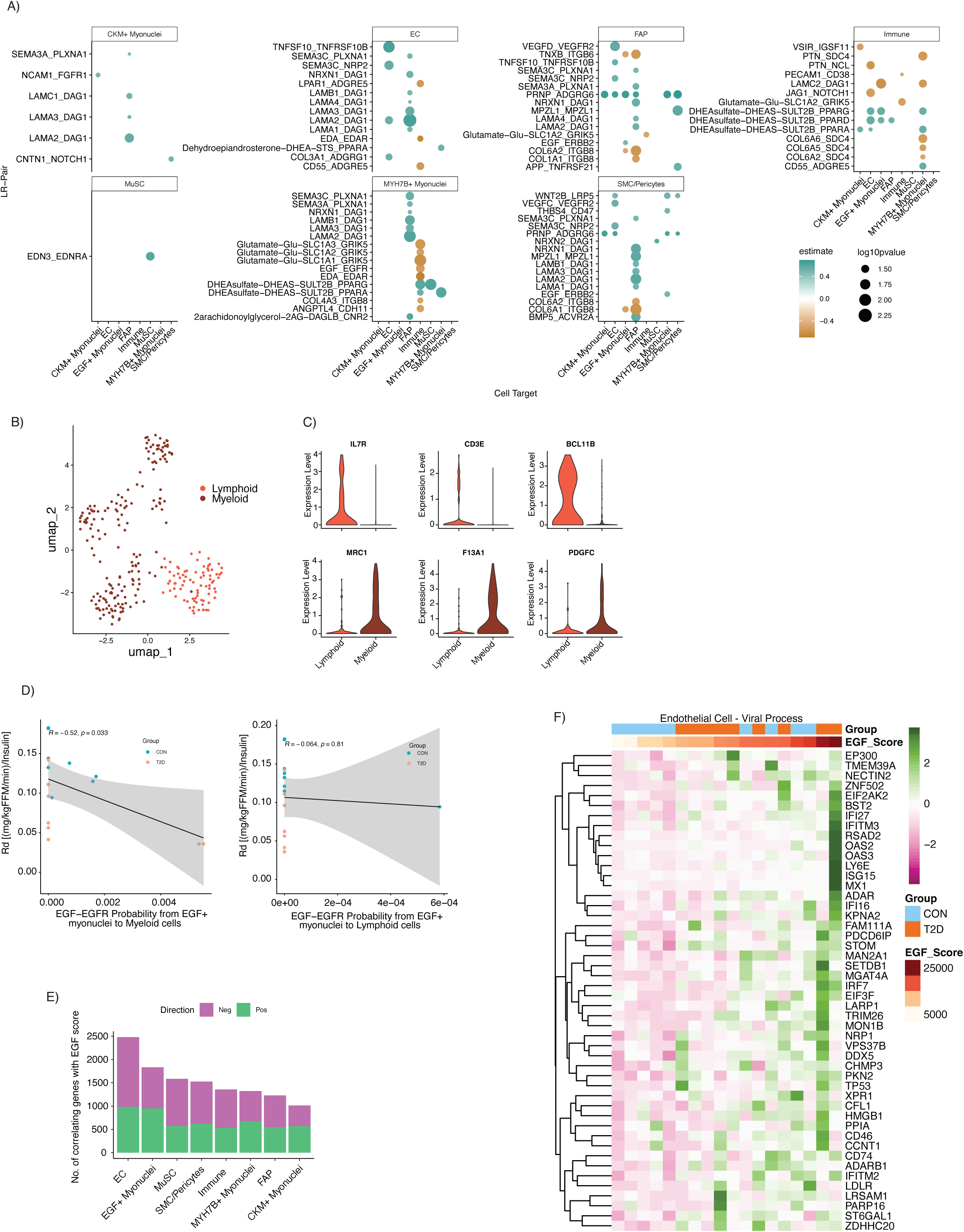
Supplement to Figure 4. Ligand-Receptor pairs that have a probability that significantly correlates with Rd/I separated by source nuclei types (A). Immune sub-clustering identified Lymphoid and Myeloid cell types (B) based on immune cell marker genes (C). EGF/EGFR ligand receptor probability from EGF+ myonuclei correlated to Rd/I when the target cell is Myeloid cells but not Lymphoid cells (D). The number of genes that positively or negatively correlate with EGF score in each nuclei sub-type (E). EGF score positively correlates with viral process genes in endothelial cells (F).

